# The effect of treatment of auxin producing bacterial culture supernatant in combination with organic and inorganic fertilizers

**DOI:** 10.1101/2022.01.31.478466

**Authors:** Seunghye Park, Ji-Hwan Shin

## Abstract

Bacteria that secret plant growth promoting substances are good candidate for field application to improve crop yield. Especially, bacteria that excrete auxin can enhance root growth and settlement of crops to help adapting new soil environment when transplanted. In this study, we applied culture supernatant of an auxin producing bacteria *Ignatzschineria* sp. CG20001. The bacterial culture supernatant (BCS) was applied in combination with chicken manure and liquid chemical fertilizer. The treatment of bacterial culture supernatant enhanced growth of red kohlrabi, *Brassica oleraceae* var. *gongyloides*, especially in combination with chicken manure. To understand mechanisms related higher growth, we compared photosynthetic efficiency by measuring chlorophyll fluorescence and light harvesting antenna pigments composition. The performance index PS II was higher in BCS treated plants, and recovered faster after leaf harvest. Photoprotective mechanism was also increased in BCS treated plants, however, and DTT infiltration, non-photochemical quenching (NPQ) reduced to similar level, implying higher xanthophyll cycle dependent NPQ induction mechanism in BCS treated plants. However, in liquid chemical fertilized plants, BCS treated and non-treated plants showed the opposite pattern in the change of NPQ. The chlorophyll *a/b* ratio was highest in no-fertilized non-treated control, and lowest in chicken manure and BCS combination, implying impact of nutrition availability between the two groups. Through these results, we conclude that the appropriate combination of organic fertilizer and plant growth promoting bacterial metabolite can improve plant growth more than chemical fertilizers. This study also provides physiological understanding for strategic fertilization regime of organic fertilizer and biofertilizer combination.

## Introduction

Due to the growing concerns for environmental problem and healthy food sources, changes in agricultural practices are drawing attention. With the growing awareness that human activities are harming the environment and nature, many efforts were made to find the sustainable solutions such as recycling and using natural resources. In agriculture, using organic fertilizers and biofertilizers are recognized as desirable practices for both environment and human health. In many cases, however, the effect of these solution did not meet the high standard set by conventional chemical fertilizers. Many efforts were made to understand and solve the low yield problems of organic fertilization (Seufert et al. 2012; Reganold and Wachter 2016; Kravchenko et al. 2017; Sharpe et al. 2020). Some advantages such as higher contents of secondary metabolites by organic fertilization seemed to come at cost of yield (Sharpe et al. 2020). The transcriptome analysis revealed the different metabolic responses by different fertilization (organic and conventional): allocation of photosynthetic products between secondary metabolites and plant growth. Agricultural practices can be more strategic through insights of metabolic changes in organic fertilized plants can improve agricultural practices in organic fertilization. However, there are gaps between the data of transcriptional level and physiological responses, due to lack of enough reports on physiological analysis.

Plant growth promoting bacteria (PGPBs) are a promising alternative to conventional agriculture, and the efficacy in crop growth is proven in many reports. They affect plant growth in direct and indirect ways, through increasing nutrient availability or suppressing pathogen growth around plants (Bernard R. Glick 1995). Production and secretion of plant hormone by PGPB is one of direct growth promoting mechanism. Many PGPBs are reported to secrete auxin, which enhanced plant root growth thereby facilitating adaptation to the soil and growth for plants, thereby enlarge ecological niche for the bacteria. This symbiotic advantage induced by bacterially produced auxin, are utilized not only plant root interacting bacteria, but also animals’ intestinal bacteria which use plants as an alternative host (Dong et al. 2003; Hernández-Reyes and Schikora 2013; Cox et al. 2018).

In spite of general agreement in the usefulness of PGPB in agriculture, the underlying mechanisms and physiological responses are not well understood. Changes in transcriptome and bacterial metabolome analysis are reported in previous studies (Sharpe et al. 2020; Morcillo et al. 2021) that explains underlying metabolic changes in transcription level, however, reports on analysis of physiological responses are rare.

Photosynthetic parameter is a prominent indicator of plants’ basic physiology whereupon all other metabolisms depends. The parameters obtained from measurement of chlorophyll fluorescence used to screen photosynthetic characteristics of crops, usually under stress conditions. Transient chlorophyll fluorescence induction measurement, O-J-I-P fluorescence measurement, offers detailed information on photosynthetic energy transfer within photosystems (Strasserf et al. 1995). A number of parameters derived from the JIP-test give structural and functional assessment on photosynthetic apparatus. The parameters are proven to be sensitive probes indicating changes in the plant’s photophysiology which is susceptible and dynamically responds to various stresses (Baker and Rosenqvist 2004), and used to study stress responses of crops in many reports (Strauss et al. 2006; Jedmowski et al. 2014; Rapacz et al. 2019).

Non-photochemical quenching (NPQ) is a mechanism that protect photosynthetic machinery from excess energy. This can be observed by measuring chlorophyll *a* fluorescence using light pulse modulation (Demmig-Adams et al. 1996). Appropriately operating NPQ is essential for maintenance of photosynthetic efficiency leading to better growth and yield.

In this study, different combination of different fertilizers and bacteria culture supernatant of *Ignatzschineria* sp. CG20001 an auxin producing bacterium (in press) was applied in crop cultivation, and compared the growth rate through repeated leaf-harvests. To understand physiological response behind the difference in growth, we analyzed photosynthetic characteristics including OJIP parameters, NPQ, and light harvesting antenna pigments composition. The results give us insight into relationships between growth rate and development of mechanisms that maintain photosynthetic efficiency under different fertilization conditions.

## Materials and methods

### Plant cultivation, fertilization and BCS treatment / leaf harvest and measurement

Seedlings of kohlrabi (*Brassica oleraceae* var. *gongylodes*) were purchased from local market, and grown in a lab under controlled environment: 90 μmol photonsom^-2^·s^-1^, 16/8 hours of light/dark cycle at 25±2°C. Plants were transferred into mixture of peat moss and vermiculite, and fertilized either with 3 grams of chicken manure compost (containing nitrogen 3%, phosphorus 2%, and potassium 1%) per pot, or one ampule of 35mL liquid chemical fertilizer (containing 0.06% nitrogen, 0.04% phosphorus, 0.03% potassium, and 0.001% manganese). The organic fertilizer was supplied once at transplantation, and the liquid chemical fertilizer was re-supplied after each leaf harvest. 1mL of BCS was supplied into each pot soil twice a week.

For preparation of BCS, *Ignatzschineria* sp. CG20001 strain were cultured in LB broth omitted NaCl. After overnighter culture, when OD600 reached about 1.0, the culture was centrifuged at 3000 rpm at room temperature for 10 minutes. The supernatant was decanted to clean containers, and kept in a refrigerator (4°C) until use.

To measure growth, leaves were harvested total four times, about eleven days intervals. At each harvest, two fully expanded youngest leaves were left. Number of harvested leaves from each plants were counted, and then collected for weighing.

### Chlorophyll fluorescence measurement

Before measuring photosynthetic parameters, leaves were covered with aluminum foil for 10 minutes for dark adaptation. Then, the photosynthetic activities were measured using FluorPen (FP100D, Photo Systems Instruments, Czech Republic). To measure NPQ, dark adapted leaves were illuminated with 500 μmol photonsom^-2^·s^-1^ of actinic light for 130 seconds, with 5 times of saturating light pulses, followed by dark relaxation for 88 seconds with 3 times of saturating light pulses. For xanthophyll cycle inhibition, 50mM of DTT solution was infiltrated into leaves, followed by 10 minutes of dark adaptation for NPQ measurement.

### Spectrophotometric pigment analysis

Five millimeter diameter of leaf discs were taken from young mature leaves. Pigments were extracted in 80% acetone. After chlorophyll were eluted in the acetone, absorbances of the supernatant were measured using spectrophotometer (UV/VIS Excellence UV5, Metler-Toledo, Switzerland). The pigment contents were calculated according to the equation below as described by Wellburn (Wellburn 1994).

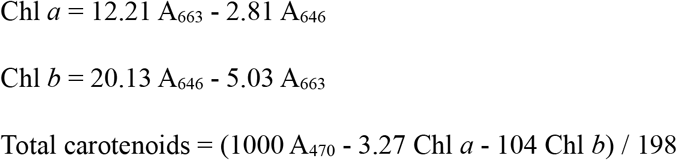

### Statistical analysis

The data are presented as the means ± standard deviations measured from three independent plants. To verify statistical significance, we performed analysis of variance (ANOVA) and Duncan’s multiple range test (DMRT) using R (4.0.2).

**Table 1.**
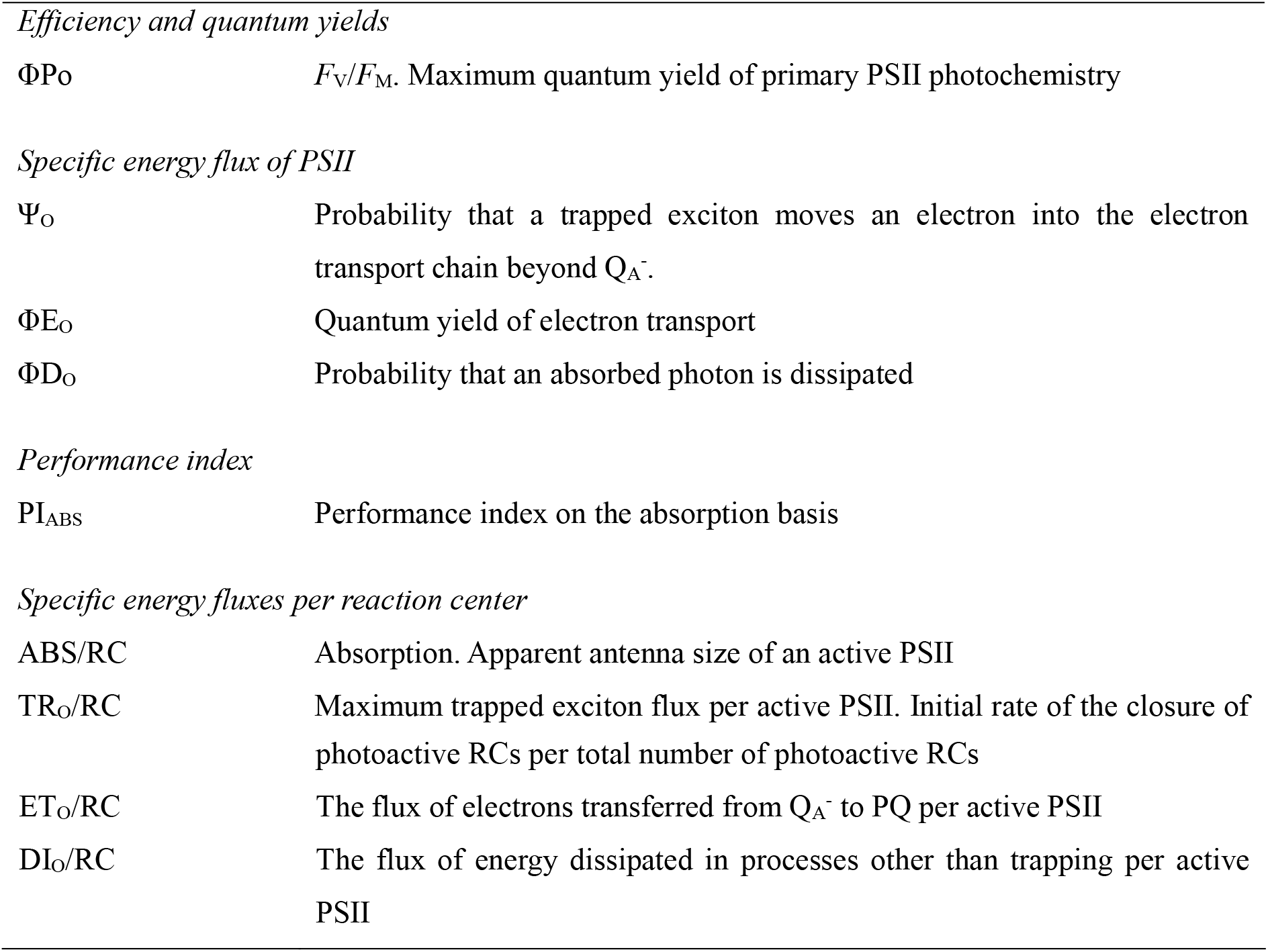
Photosynthetic parameters obtained from transient chlorophyll fluorescence (Strasser et al. 2000).

|  |  |
| --- | --- |
| *Efficiency and quantum yields* |  |
| $\Phi_{Po}$ | $F_v/F_m$. Maximum quantum yield of primary PSII photochemistry |
| *Specific energy flux of PSII* |  |
| $\Psi_o$ | Probability that a trapped exciton moves an electron into the electron transport chain beyond $Q_A^-$. |
| $\Phi_{Eo}$ | Quantum yield of electron transport |
| $\Phi_{Do}$ | Probability that an absorbed photon is dissipated |
| *Performance index* |  |
| $PI_{ABS}$ | Performance index on the absorption basis |
| *Specific energy fluxes per reaction center* |  |
| ABS/RC | Absorption. Apparent antenna size of an active PSII |
| TR<sub>o</sub>/RC | Maximum trapped exciton flux per active PSII. Initial rate of the closure of |
|  | photoactive RCs per total number of photoactive RCs |
| ET<sub>O</sub>/RC | The flux of electrons transferred from Q<sub>A</sub><sup>-</sup> to PQ per active PSII |
| DI<sub>O</sub>/RC | The flux of energy dissipated in processes other than trapping per active PSII |

## Results

To understand the combinatorial effects of different kinds of fertilizers and CG20001 bacterial culture supernatant (BCS), we applied BCS to three differently fertilized plants: chicken manure (as an organic fertilizer), liquid chemical fertilizer, and no-fertilized control, and compared the effects with non-treated plants (Fig. 1A).

**Fig. 1.**
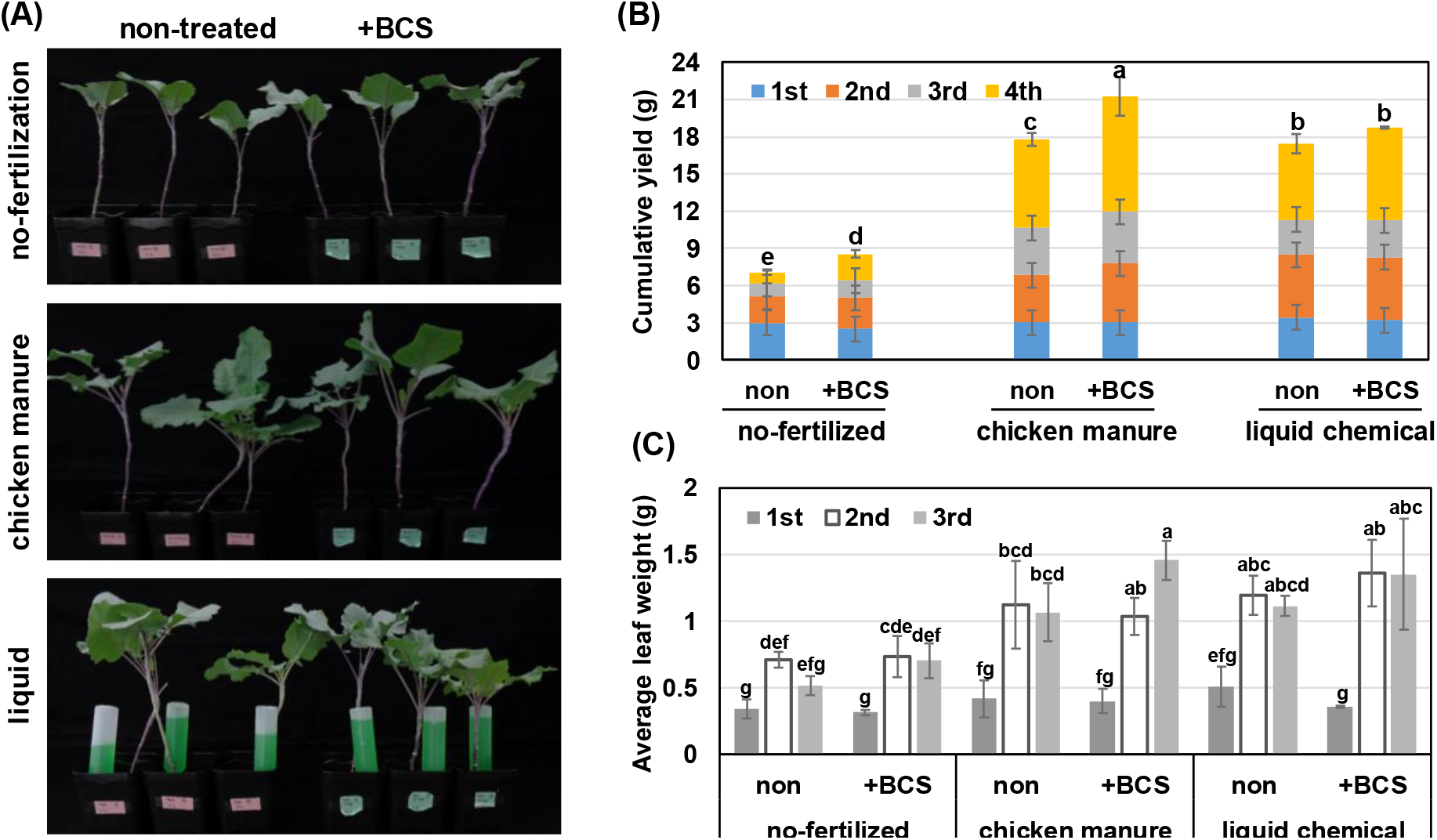
Experimental setting and growth measurement of red kohlrabi (*Brassica oleraceae* var. *gongylodes*). (A) Pot grown red kohlrabi plants right before the third leaf growth measurement. (B) Cumulative leaf mass through the four times of harvests. (C) Average weight of harvested leaves in the first three harvests. Error bars represent standard deviation of three independent plants. Significantly different treatment groups (*p*<0.05) are represented with different letters over each bar.

### Plant growth and leaf mass production

Vegetative growth of differently fertilized plants were measured from fresh leaf weight. Fertilization greatly improved leaf production, which was about two to three times higher than non-fertilized control (Table 2). The cumulative mass was the greatest in the BCS treated plants in combination with chicken manure: the mass was about three times higher (21.26±2.22 g) than that of non-fertilized no-treated control (6.95±0.33 g). BCS treatment slightly increased leaf mass in liquid chemical fertilization and non-fertilized control plants, however, the differences were not statistically significant among liquid chemical fertilized plants. Without fertilization, treatment of BCS still produced about 16% more leaf mass than the non-treated control plants (Fig. 1B).

**Table 2.**
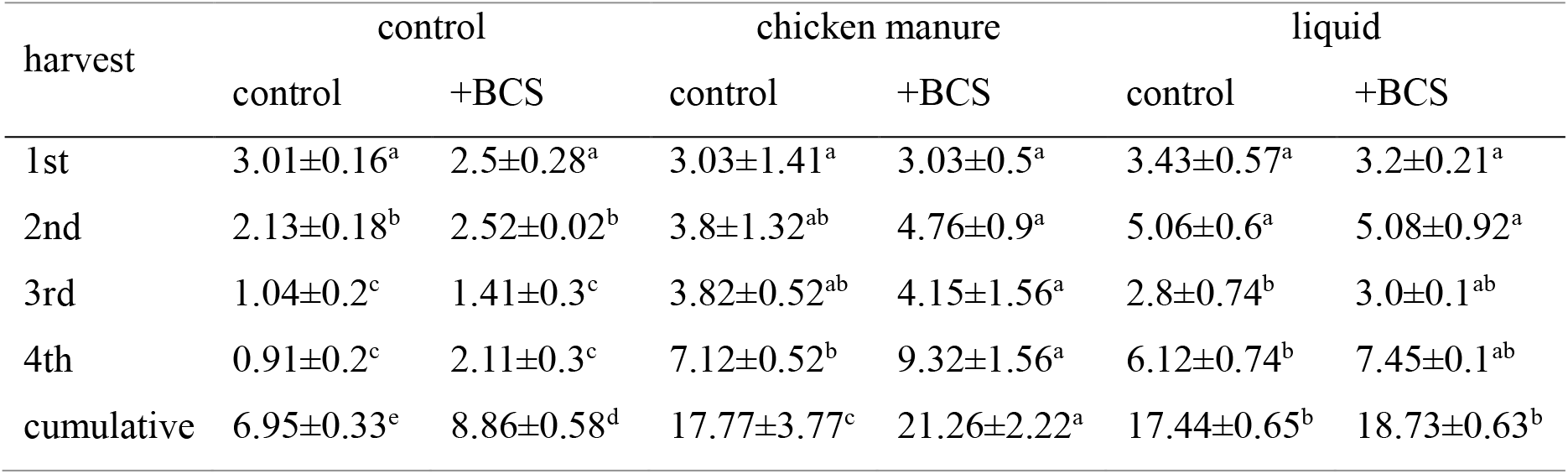
Harvested mass from differently fertilized *B.oleracea* var. gongylodes plants through four times of leaf harvest. Values represent average fresh weight of leaves harvested from three independent plants with standard deviation. Means with the same letter are not significantly different (*p*>0.05).

The sizes of leaves also differed in different fertilization, to compare size of leaves as weight, harvested masses were divided with numbers of harvested leaves. The average weights of leaf in chicken manure and liquid chemical fertilized plants were heavier than no-fertilized control. BCS treatment did not affect leaf weight (or size), except for chicken manure fertilized plants. In chicken manure +BCS combination, leaf weight increased at each harvest, and in the third harvest, the plants produced the biggest leaves (1.46±0.14 g), compared to chicken manure fertilized plants without BCS treatment (1.07±0.2 g) (Fig. 1C).

### Photosynthetic parameters

We hypothesized that the higher yields in the BCS treated plants, stems from better photosynthetic activities. To assess photosynthetic properties, we measured transient chlorophyll fluorescence induction, known as OJIP transients, between the third and the forth harvest. During the three weeks of measurement, the performance index parameter, PI_ABS_, changed most dynamically. The parameter decreased after harvest, and increased again until 23th days after harvest, and recovered faster until 23th days after harvest in BCS treated plants, however in BCS non-treated plants, only liquid chemical fertilized plants recovered. Other parameters known to be responding to stress, such as ETo/RC, DIo/RC changed only slightly in the during the eleven days after harvest, and *F*_V_/*F*_M_ (ΦPo) did not change significantly (Fig. 2).

**Fig. 2.**
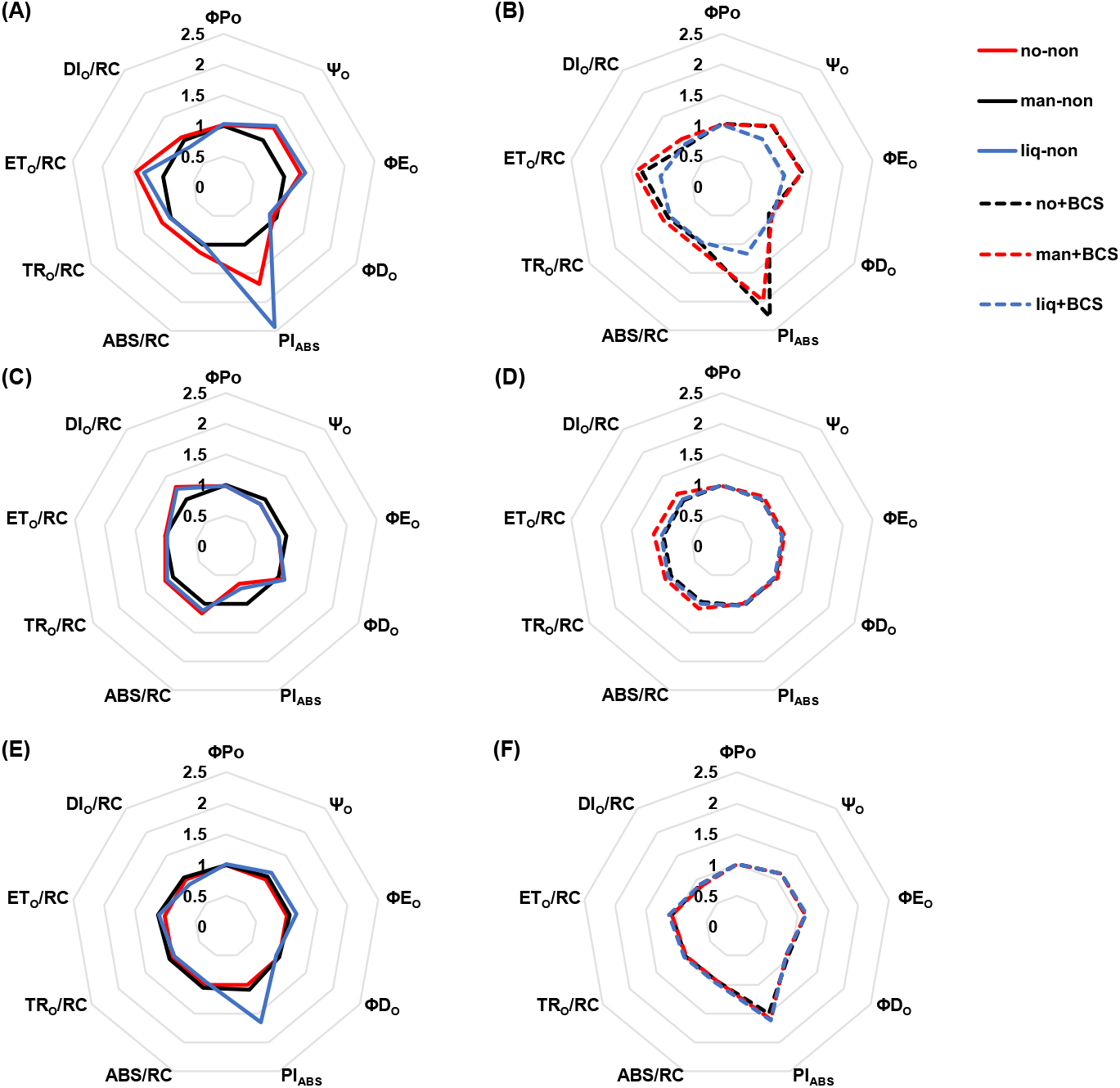
Photosynthetic properties measured through transient chlorophyll fluorescence, OJIP parameters. Leaves of plants grown in different fertilization regime: control, chicken manure, and liquid fertilizer with (B, D, F) or without (A, C, E) BCS. After the third harvest, OJIP test were performed three times, right after the third harvest (A), (B), 11 days later (C), (D), and 23 days after harvest (E), (F). values are average obtained from measurement of three independent plants, and normalized to the control sample.

### Non-photochemical quenching induction

For development and maintenance of photosynthetic efficiency, photoprotection is of critical importance. To compare the photoprotective capacity, NPQ was measured in normal leaves and DL-dithiothreitol (DTT) treated leaves. DTT inhibits zeaxanthin deepoxidase activity, thereby blocks xanthophyll cycle which is one of the fastest induced NPQ components. By comparing NPQ induction of DTT treated samples with untreated samples, we can roughly estimate xanthophyll cycle dependence in NPQ of the particular samples.

In our measurement, NPQ induction was higher in BCS treated plants among chicken manure fertilized plants and non-fertilized plants. On the contrary, in liquid chemical fertilized plants, higher NPQ was induced in non-treated plants on average, however, the difference was not significant when the variations among the plants were taken into account (Fig. 3).

**Fig. 3.**
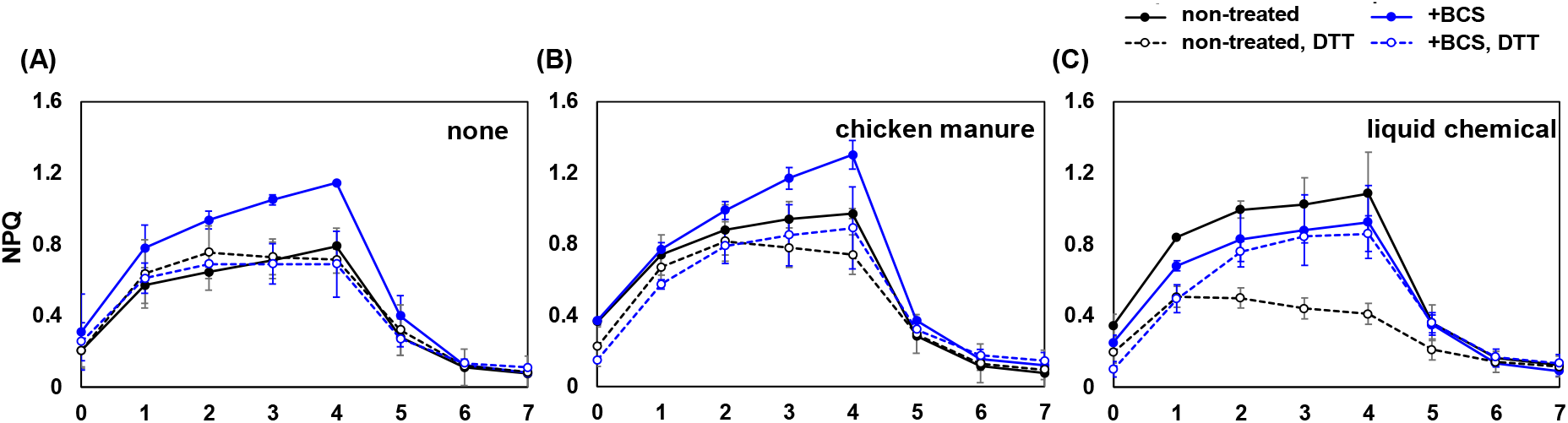
NPQ induction pattern of red kohlrabi grown under different fertilization scheme: (A) non-fertilized, (B) chicken manure, and (C) liquid chemical fertilized plants. Dashed lines indicate NPQ induction of DTT treated plants, and solid lines non treated plants. Black color are BCS-non-treated plants, and blue colored lines are BCS treated plants. NPQ are represented with standard deviation measured from three independent plants.

When DTT was infiltrated to the leaves, level of NPQ slightly reduced in BCS non-treated plants in no-fertilized and chicken manure fertilized plants. However, the reduction in BCS treated plants was more pronounced in these fertilization. On the contrary, in liquid chemical fertilized plants, DTT infiltration brought the opposite effect, and distinctive reduction of NPQ in BCS non-treated leaves was observed.

### Photosynthetic pigment composition

To compare light harvesting antenna pigment composition, Chlorophyll *a*/*b* ratio and carotenoid to chlorophyll ratio of leaves were analyzed. Each fertilization group showed significant difference in Chl *a*/*b* ratio, between BCS treated and non-treated. Overall, Chl *a*/*b* ratio was higher in BCS non-treated compared to BCS treated. The Chl *a*/*b* ratio was the highest in no-fertilized non-treated control, and the lowest in chicken manure fertilized in combination with BCS treatment. The liquid chemical fertilized plants showed little difference (Fig. 4A).

**Fig. 4.**
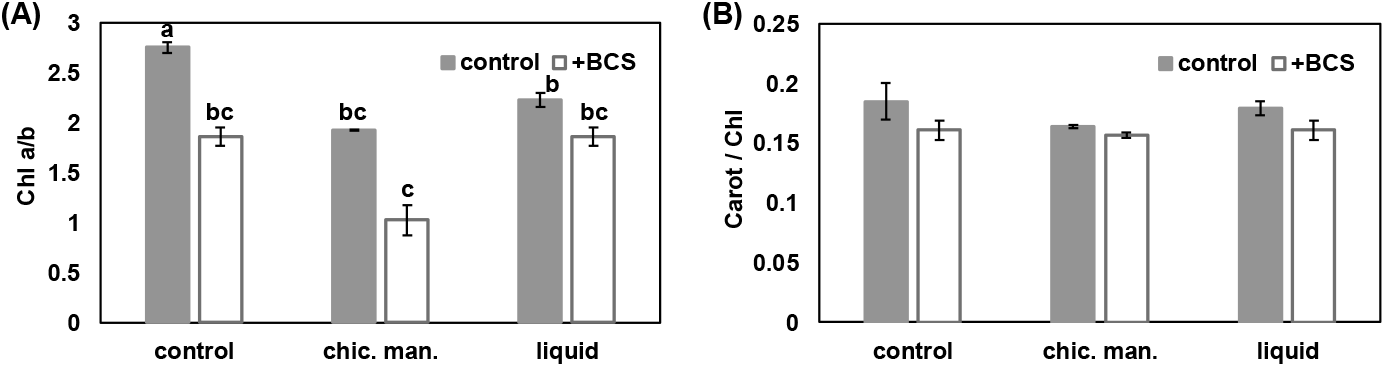
Photosynthetic pigment composition in leaves of differently fertilized kohlrabi. (A) Ratio of Chl *a* to *b*. (B) Ratio of total carotenoid to total chlorophyll. The treatments with the same letters on the column are not significantly different at 0.05 probability level.

On the contrary, no significant difference was found in carotenoid to Chl ratio. The average carotenoid to Chl ratio was slightly higher in BCS non-treated plants in no-fertilization and liquid chemical fertilized plants, however, statistically there was no significant difference (Fig. 4B).

## Discussion

The mechanism behind the effects of PGPB and organic fertilization is hard to understand, because, unlike inorganic fertilizers they include various unidentified effectors that might act through indirect ways of affecting plant physiology. Nutritious impact of organic fertilizers can be improved by combinatorial inoculation of PGPB. Due to increase in nutrient availability by PGPB activity, crop yield could be increased in combination with organic fertilizers (Suliasih and Widawati 2018). IAA produced by the bacteria in those studies supposed to help growth and activity of plant roots. In this study, IAA containing BCS, without bacterial cells, improved plant growth in combination with chicken manure, an organic fertilizer. Considering little difference of yield in liquid chemical fertilized plants, we could speculate that bacterial metabolite synergistically affected plant growth.

Analyzing photosynthetic activity on which plant growth depends, gave us an insight into the physiology behind better growth. OJIP parameters are known to sensitively respond to stresses, and PI_ABS_ is the best indicator of photosystem functionality (Strauss et al. 2006). In this study, PI_ABS_ is the parameter that showed most distinctive change with time between the harvest terms among different fertilization and treatment. Drastic reduction of PI_ABS_, known to represent plant vitality (R.J. Strasser, A. Srivastava 2000), indicate plants are experiencing stress, and more severely in non-treated plants compared to BCS treated. The most distinctive characteristics of the bacteria CG20001 was auxin production, and its culture supernatant used in this study contained about 50ppm of IAA. There is no direct link of auxin regulating photosynthetic activity, however, auxin treatments are reported to influence photosynthetic activity through indirect ways. Hayat et al. (Hayat et al. 2009) reported treatment of auxins and the analogues improved nitrogen metabolism and photosynthesis. The faster recover of PI_ABS_ in the BCS treated plants in this study might be in line with their observation.

Other parameters did not change significantly. It is probably, that the stress conditions where OJIP parameters changed substantially in other reports, the photosynthetic apparatus also had undergone significant changes. The maximum quantum yield, ΦPo (=*F*_V_/*F*_M_), reportedly does not change untill the plant undergoes severe stress beyond recover, such as loss of 90% loss of chlorophylls (Oukarroum et al. 2009; Osório et al. 2014; Manaa et al. 2021). Changes of other parameters, for example, slight increase in dissipation parameter (ΦDo, DIo/RC) and inactivation parameter of PSII reaction center (ABS/RC) was observed in nutrient-deficient leaves in previous report (Kalaji et al. 2014).

In order to maintain photosynthetic efficiency, it is important to have operating photoprotective mechanisms. Plants have developed many mechanisms that adjust photosynthetic apparatus to environmental conditions and physiological balance. Light harvesting antenna proteins are essential in photosynthesis and susceptible to photodamage that requires frequent repair, therefore, synthesis and maintenance of them are burden to plants under nutrient limitation. The increase of Chl *a/b* ratio usually are incurred by reduction of Chl *b* synthesis resulting in reduction in total Chl contents and size of light harvesting antenna complexes (LHCs). Nitrogen availability alongside high light intensity have been reported to affect this change (KITAJIMA and HOGAN 2003). In addition to nitrogen, other nutrients also affected Chl biosynthesis and photosynthetic energy flow (Abadía 1992; Ciompi et al. 1996; D’Hooghe et al. 2013). In that context, the highest Chl *a/b* ratio in no-fertilized non-treated control and the lowest in chicken manure and BCS treatment in our study, might be resulted from different nutritional conditions.

The reduction in the amount of Chl *b* is a part of photoprotective mechanism, however, the presence of Chl *b* is essential for stability of light harvesting antenna complexes. Therefore, in addition to reduction in Chl *b* content, additional photoprotective mechanisms are required working in different mode of action. NPQ is a photoprotective mechanism that can be measured through Chl *a* fluorescence using pulse amplitude modulating (PAM) devices that provides parameters representing dynamic kinetics of light energy flow through photosynthetic electron transport chains (Roháček 2002; Baker 2008). The high induction of NPQ represent appropriately operating NPQ in BCS-treated, especially in combination with chicken manure fertilized plants in this study. Photoprotective mechanisms which seemed as inhibitor of efficient photosynthesis in the past (Müller et al. 2001), are proven to be the critical contributor enabling harmless and balanced photosynthesis in the fluctuating environment (Wu et al. 2020).

One of the four components of NPQ is qZ which depends on xanthophyll cycle that mediated by zeaxanthin deepoxidase (Latowski et al. 2011). Treating DTT, the inhibitor of zeaxanthin deepoxidase, the proportion of qZ in NPQ can be estimated through reduction of NPQ. It is not clear if the different degree of NPQ reduction between liquid chemical fertilized and the other groups (by DTT treatment) are actually resulted from different dependence on xanthophyll cycle, until precise pigment compositions are estimated with high-resolution analysis.

It seems that there is no direct relation between NPQ and external auxin treatment. However, NPQ and photosynthetic energy flow is affected by nutritional factors. Su et al. (Su et al. 2020) reported change of photosynthetic energy flow affected by zinc (Zn) availability that might be influenced by auxin. Increased aluminum (Al) solubility in acidic soil disturbed IAA biosynthesis through competition with Zn which is essential component of IAA biosynthesis, however, the Al effect was complemented by supply of Zn. The addition of Zn caused down-regulation of CEF-dependent generation of proton gradient (ΔpH_*pmf*_), thereby, reduced oxidized state of Q_A_ and reaction centers of photosystems, increasing ability of photosystems. Therefore, in that case, IAA was in the hub of Zn absorption and detrimental effect of Al, affecting photosynthetic energy flow.

The xanthophyll cycle is activated by proton gradient across thylakoid membrane, which is related to another NPQ component, qE (Li et al. 2002). This NPQ component is not dependent of xanthophyll pigment, however, closely related to the condition that induces qZ. Therefore, different nutritional condition in this study might have caused induction of NPQ in different ways.

Comprehensively, photosynthetic performance and photoprotective mechanism are most distinctively activated in chicken manure fertilized in combination with BCS treated plants. This effect might be combinatorial effects of nutrient supplied by chicken manure and the IAA contained in the BCS. However, there might be other factors that enhance their effects, considering the synergy effect was not observed in the liquid chemical fertilization in spite of comparable nutrient composition. Therefore, it requires further analysis of the BCS, physiological aspects of the treated plants.

## Conclusion

In this study, we compared different crop growth and physiology, treated with different fertilizers with BCS combination. The growth measured by leaf mass production, was 13.5% higher in chicken manure + BCS combination than chemical + BCS combination. The difference between BCS treated and non-treated was 2.7 times higher in chicken manure fertilization than liquid chemical fertilization. The photosynthetic parameters indicate that better growth of chicken manure fertilized in combination with auxin producing bacteria culture supernatant, is partly related to efficient photosynthesis and photoprotection. To generalize the synergy effects, more organic fertilizers need to be tested. However, lack of the such effects observed in chemical fertilization, support the idea of synergy effect of BCS with organic fertilization. For the bacterial strain used in this study, need to be analyzed for its nutrient solubilizing ability, and presence of other effectors. It is desirable to replace chemical fertilizers with organic and biofertilizers. More detailed studies based on the observation in this study, will be useful in finding out optimal combinations for specific crops, and estimation of their effects.

